# Contextual diversity of the human cell-essential proteome

**DOI:** 10.1101/107797

**Authors:** Thierry Bertomeu, Jasmin Coulombe-Huntington, Andrew Chatr-Aryamontri, Karine Bourdages, Yu Xia, Mike Tyers

**Author notes:** Co-first authors (Thierry Bertomeu, Jasmin Coulombe-Huntington, Andrew Chatr-Aryamontri). Abbreviations: universal essential (UE), contextual essential (CE), lone essential (LE), non-essential (NE), single-guide RNA (sgRNA).

## Abstract

Essential genes define central biological functions required for cell growth, proliferation and survival, but the nature of gene essentiality across human cell types is not well understood. We assessed essential gene function in a Cas9-inducible human B-cell lymphoma cell line using an extended knockout (EKO) library of 278,754 sgRNAs that targeted 19,084 RefSeq genes, 20,852 alternatively-spliced exons and 3,872 hypothetical genes. A new statistical analysis tool called RANKS identified 2,280 essential genes, 234 of which had not been reported previously. Essential genes exhibited a bimodal distribution across 10 cell lines screened in different studies, consistent with a continuous variation in essentiality as a function of cell type. Genes essential in more lines were associated with more severe fitness defects and encoded the evolutionarily conserved structural cores of protein complexes. Genes essential in fewer lines tended to form context-specific modules and encode subunits at the periphery of essential complexes. The essentiality of individual protein residues across the proteome correlated with evolutionary conservation, structural burial, modular domains, and protein interaction interfaces. Many alternatively-spliced exons in essential genes were dispensable and tended to encode disordered regions. We also detected a significant fitness defect for 44 newly evolved hypothetical reading frames. These results illuminate the nature and evolution of essential gene functions in human cells.

## Introduction

Essential genes underpin the genetic architecture and evolution of biological systems (Giaever et al. 2002). In all organisms, essential genes are needed for survival and proliferation, while in multicellular organisms additional essential genes function in different tissues at various stages of development. In the budding yeast *Saccharomyces cerevisiae*, only 1,114 of the ~6,000 genes encoded in the genome are essential for growth in nutrient-rich conditions (Giaever et al. 2002). The non-essential nature of most genes suggests that the genetic landscape of the cell is shaped by redundant gene functions (Hartman et al. 2001). Consistently, systematic screens in *S. cerevisiae* have uncovered more than 500,000 binary synthetic lethal interactions (Tong et al. 2001; Costanzo et al. 2010; Costanzo et al. 2016). In parallel, context-specific chemical screens have revealed that virtually every gene can be rendered essential under the appropriate condition (Hillenmeyer et al. 2008). The prevalence of non-essential genes has been verified by systematic genetic analysis in other single-celled organisms including *Schizosaccharomyces pombe* (Kim et al. 2010), *Candida albicans* (Roemer et al. 2003) and *Escherichia coli* (Baba et al. 2006). In metazoans, the knockdown of gene function by RNAi in the nematode worm *Caenorhabditis elegans* revealed that 1,170 genes are essential for development (Kamath et al. 2003), while in the fruit fly *Drosophila melanogaster* at least 438 genes are required for cell proliferation in vitro (Boutros et al. 2004). In the mouse *Mus musculus*, ~25% of genes tested to date are required for embryonic viability (Dickinson et al. 2016). Essential genes tend to be highly conserved, to interact with one another in local modules, and be highly connected in protein and genetic interaction networks (Hirsh and Fraser 2001; Zotenko et al. 2008; Costanzo et al. 2016). Although essentiality is often framed as an all-or-none binary phenotype, in reality the loss of gene function causes a spectrum of fitness defects that depend on developmental and environmental contexts. For example, in *S. cerevisiae* an additional ~600 genes are required for optimal growth in rich medium (Giaever et al. 2002) and in *C. elegans* most genes are required for fitness at the whole organism level (Ramani et al. 2012). The definition of essentiality is thus dependent on context and the experimental definition of fitness thresholds.

The clustered regularly interspaced short palindromic repeats (CRISPR) and CRISPR-associated (CAS) protein system directs the cleavage of specific DNA sequences in prokaryotes, where it serves as an adaptive immunity mechanism against infection by foreign phage DNA (Samson et al. 2013). The Cas9 endonuclease can be targeted toward a specific DNA sequence by a single-guide RNA (sgRNA) that contains a 20 nucleotide match to the locus of interest (Jinek et al. 2012). The co-expression of Cas9 and an sgRNA leads to a blunt end double-strand break (DSB) at a precisely specified position in the genome. Repair of the DSB by error-prone non-homologous end joining (NHEJ) leads to random DNA insertions or deletions (indels) that cause frame-shift mutations and an ersatz knockout of the target gene with high efficiency (Cong et al. 2013).

CRISPR/Cas9 technology has been recently adapted to perform large-scale functional gene knockout screens in human cells (Shalem et al. 2015). A pooled library of 64,751 sgRNAs that targets 18,080 genes was used in screens against melanoma and stem cell lines (Sanjana et al. 2014; Shalem et al. 2014), while a library of 182,134 sgRNAs that targets 18,166 genes was used in screens against chronic myelogenous leukemia and Burkitt’s lymphoma cell lines (Wang et al. 2015). A third library of 176,500 sgRNAs that targets 17,661 genes was used in screens against colon cancer, cervical cancer, glioblastoma and hTERT-immortalized retinal epithelial cell lines (Hart et al. 2015). In parallel, genome-scale screens based on transposon-mediated gene-trap technology have been performed in two haploid human cell lines (Blomen et al. 2015; Wang et al. 2015). Each of these genome-wide screens identified on the order of 1,500-2,200 essential genes that were enriched in functions for metabolism, DNA replication, transcription, splicing and protein synthesis (Blomen et al. 2015; Hart et al. 2015; Wang et al. 2015). Importantly, the CRISPR/Cas9 and gene-trap screens exhibit a strikingly high degree of overlap and overcome the limitations of previous RNAi-based gene knockdown approaches (Blomen et al. 2015; Hart et al. 2015; Wang et al. 2015).

Currently available human genome-wide CRISPR libraries target most well-characterized gene loci. Two libraries were designed to target the well-validated RefSeq gene collection (Sanjana et al. 2014; Wang et al. 2015) while a third library targets confirmed protein coding gene regions in the GENCODE v17 assembly (Hart et al. 2015). To extend functional CRISPR/Cas9 screens to less well characterized regions of the genome at potential sub-gene resolution, we generated a high complexity extended knockout (EKO) library of 278,754 sgRNAs that targets 19,084 RefSeq genes, 20,852 unique alternative exons and 3,872 hypothetical genes. The EKO library also includes 2,043 control sgRNAs with no match to the human genome to estimate screen noise. We used the EKO library to identify essential genes in the NALM-6 human pre-B cell lymphocytic leukemia line, including alternatively-spliced exons and hypothetical genes not previously tested in CRISPR/Cas9 screens. Our analysis revealed the broad influence of Cas9-induced double-stranded breaks on cell viability, the residue-level determinants of essential protein function, the prevalence of non-essential exons, and the evolution of new essential genes. Integration of our data with nine previous genome-wide CRISPR/Cas9 screens revealed a bimodal distribution of essential genes across cell lineages and a distribution of subunit essentiality across protein complexes, consistent with a continuous distribution of gene essentiality. High resolution CRISPR/Cas9 genetic screens can thus uncover the organization and evolution of the essential human proteome.

## Results

### Design and generation of the EKO library

We generated a custom sgRNA library that targeted more genomic loci than previously published libraries, including additional RefSeq genes, alternatively-spliced exons and hypothetical protein-coding genes (Fig. 1A; see Table S1 for all sgRNA sequences). The core set of the EKO library was comprised of 233,268 sgRNAs that target 19,084 RefSeq protein-coding genes and 17,139 alternatively-spliced exons within these coding regions. An extended set of 43,443 sgRNAs within the EKO library was designed to target loci predicted to be potential protein coding regions by AceView (Thierry-Mieg and Thierry-Mieg 2006) or GENCODE (Harrow et al. 2012), as well as additional alternatively-spliced exons predicted by AceView that we validated by independent RNA-seq evidence. Each gene in the EKO library was targeted by approximately 10 sgRNAs and each alternative exon by 3 sgRNAs. The combined gene list represented in the EKO library contains almost 5,000 more candidate genes than any sgRNA library screened to date, including almost 1,000 additional RefSeq genes (Sanjana et al. 2014; Shalem et al. 2014; Hart et al. 2015; Wang et al. 2015). To estimate the effect of noise in our screens, we also designed 2,043 sgRNAs that had no sequence match to the human genome.

**Figure 1.**
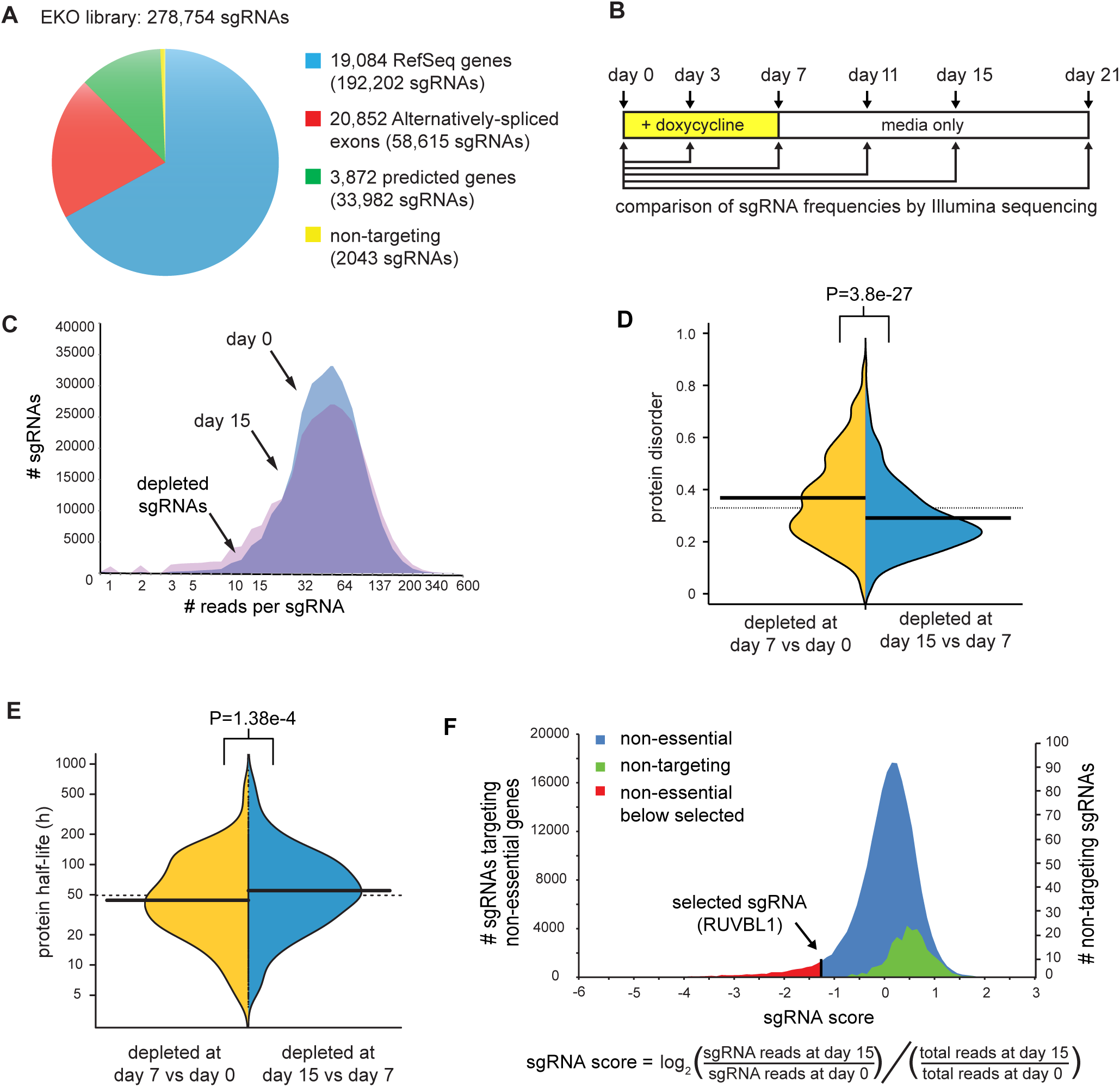
Genome-wide sgRNA library generation and screen for essential genes. **(A)** Composition of the EKO library. Some sgRNAs were used to evaluate both an alternative exon and the entire gene. **(B)** Experimental design for screen. A NALM-6 doxycycline-inducible Cas9 cell line was transduced with the EKO library at MOI of ~0.5, followed by sgRNA vector selection, population outgrowth and determination of sgRNA frequencies at the indicated time points. **(C)** Number of sgRNA reads pre-doxycycline induction (day 0) superimposed on distributions after doxycycline induction (day 7) and a further 8 days outgrowth in media only (day 15). A constant factor was used to center the two distributions. **(D)** Effect of protein intrinsic disorder on kinetics of sgRNA depletion. Average probability of disordered stretches over the entire protein as predicted by IUPred (Dosztanyi et al. 2005) for the 1,000 genes with most highly depleted sgRNAs at day 7 versus versus day 15, two-tailed Wilcoxon test. **(E)** Effect of target protein half-life on kinetics of sgRNA depletion. Half-life of mouse orthologs of the 1,000 most highly depleted genes after day 7 (N=645) was compared to day 15 (N=724). P-value, two-tailed Wilcoxon test. **(F)** Distribution of log_2_ sgRNA frequency changes for day 0 versus day 15 for all previously described non-essential sgRNAs (Blomen et al. 2015; Hart et al. 2015; Wang et al. 2015) compared to non-targeting sgRNAs in this study. An sgRNA targeting an essential gene in this study (RUVBL1) is shown for reference. Single-tailed p-values for each sgRNA were calculated as the area under the curve of the control distribution to the left of the sgRNA score divided by the total area under the control curve.

**Table 1.** Multi-variate Pearson correlation between sgRNA features and gene-normalized sgRNA depletion score.

| sgRNA feature | p-value |
| --- | --- |
| Overlapping domain | 0.0018 |
| Relative conservation | 1.33e-6 |
| Relative burial | 3.23e-6 |
| Overlapping alpha-helix | 8.27e-5 |
| Overlapping beta-strand | 8.33e-6 |
| Overlapping PPI interface | 0.0034 |
| Overlapping protein disorder | 3.30e-5 |

### Identification of cell fitness genes with the EKO library

We used the EKO library in a viability screen to assess fitness defects caused by loss of gene function in the human pre-B cell acute lymphoblastic leukemia NALM-6 line, which is pseudodiploid and grows rapidly in suspension culture (Hurwitz et al. 1979). A tightly regulated doxycycline-inducible Cas9 clone of NALM-6 for use in library screens was generated by random integration of a lentiviral construct. The EKO library was transduced at low multiplicity of infection (MOI) and selected for lentiviral integration on blasticidin for 6 days, after which Cas9 expression was induced for 7 days with doxycycline, followed by outgrowth for 15 more days in the absence of doxycycline (Fig. 1B). Changes in sgRNA frequencies across the entire library were monitored by Illumina sequencing at six different time points during the screen. The EKO library pool was well represented after blasticidin selection (i.e., the day 0 time point for the screen), with 94% of all sgRNA read counts falling within a 10-fold range relative to one another (Fig. 1C; Fig. S1A). Little change in sgRNA frequency was observed after only 3 days of doxycycline induction (Fig. S1B), likely due to the lag in Cas9 induction, the kinetics of indel generation, and the time required for effective protein depletion. A progressively larger spread in sgRNA read frequencies was observed over the 21 day time course (Fig. S1C-F). We observed that genes with greater sgRNA depletion at earlier time points tended to encode proteins that were more disordered (Fig. 1D) and that had a shorter half-life (Fig. 1E). However, genes depleted by day 21 did not possess significantly longer half-lives than those depleted by day 15 (p>0.05, Wilcoxon test, data not shown), and genes that scored in the top 2,000 most depleted at day 21 but not at day 15 sample were less likely to be validated by gene-trap scores in the HAP1 cell line (p=2.37e-6, Wilcoxon test; Fig. S1G). Based on these results, we chose to use the sgRNA read frequencies at day 15 for our subsequent analyses, with the day 0 frequencies serving as the reference time point.

### Calculation of significant sgRNA depletion by RANKS

To compute the relative level of depletion of each sgRNA, we developed a custom tool called Robust Analytics and Normalization for Knockout Screens (RANKS), which enables statistical analysis of any pooled CRISPR/Cas9 library screen or shRNA screen. First, the log_2_-ratio of the sgRNA read frequency at the day 15 versus day 0 time points, normalized by the total read count ratio, was used to quantify the relative abundance of each sgRNA (Fig. 1G). The average log_2_-ratio for all ~10 sgRNAs that target each gene was used to generate a gene score (Wang et al. 2015), which reflected the potential fitness defect for each gene knockout. To account for experimental variation in sgRNA read counts within any given screen, instead of using the log_2_-ratio of the sgRNA itself, we used RANKS to estimate a p-value based on how each log_2_-ratio compared to a selected set of non-targeting control sgRNAs. The resulting gene log p-value score was then obtained from the average of the log p-values for each of the 10 sgRNAs per gene. As applied to our dataset, RANKS performed better than previous other methods (Li et al. 2014; Hart et al. 2015; Wang et al. 2015) as judged by established correlates (Fig. S2; Table S2). Subsequent analyses were based on the RANKS statistical scores for each gene, exon or hypothetical gene represented in the EKO library.

### Genome-scale correlates of sgRNA depletion

The results of the NALM-6 screen revealed that fitness scores correlated well with features known to be associated with essential genes. We examined correlates with mutation rate as estimated by protein sequence conservation across 46 vertebrate species (Blanchette et al. 2004), node degree in protein-protein interaction (PPI) networks (Chatr-Aryamontri et al. 2015), mRNA expression level in NALM-6 cells, essentiality detected by the gene-trap method (Blomen et al. 2015), essentiality detected by a whole-genome CRISPR/Cas9 screen in the near haploid KBM7 cell line (Wang et al. 2015), and DNA accessibility as assessed by DNAse1 hypersensitivity peak density in naïve B cells (Fig. 2A; Fig. S3). For every feature, we observed a highly significant difference between the top-ranked 2,000 genes in our screen (1-2000) and the next 2,000 ranked genes (2,001-4,000), with p-values <0.05 (Wilcoxon test), such that each feature correlated strongly with gene essentiality. The strong agreement between depletion scores from an independent genome-wide CRISPR screen with our ranked gene list over the entire distribution confirmed the reproducibility of the CRISPR/Cas9 method.

**Figure 2.**
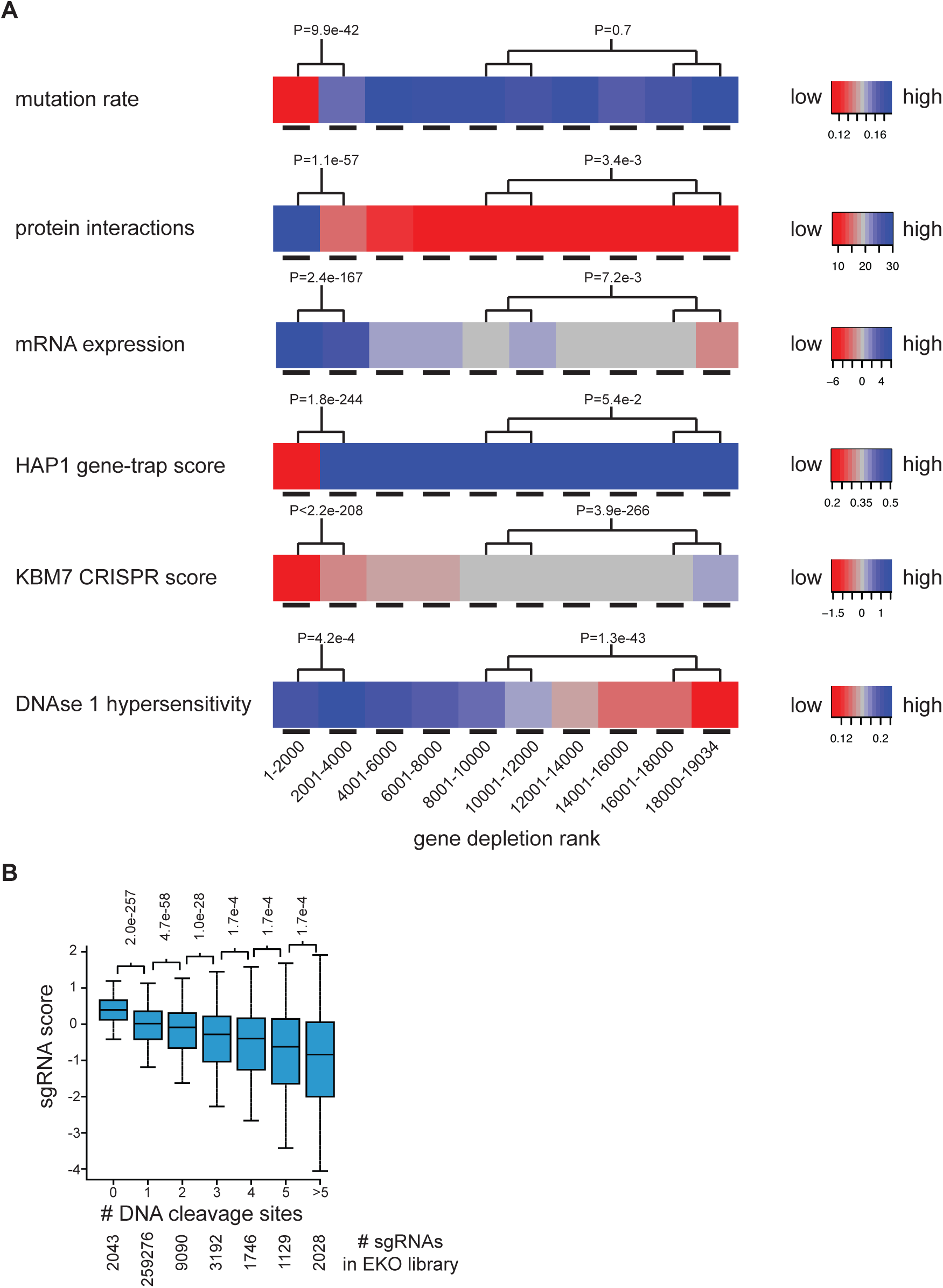
Features correlated with gene depletion. **(A)** RefSeq genes were ranked in bins of 2,000 genes from most to least depleted sgRNAs. Mutation rate was the fraction of aligned residues that differed from the human sequence across 45 vertebrate species in a 46-way Multi-Z whole-genome alignment. Protein interaction was the number of partners reported in the BioGRID database (3.4.133 release). mRNA expression was the log_2_ RNA-seq reads per million (RPM) in the NALM-6 cell line. HAP1 gene-trap score was the ratio of sense to anti-sense intronic insertions (Blomen et al. 2015). KBM7 CRISPR score was the average log_2_ read frequency change (Wang et al. 2015). DNAse1 hypersensitivity was the number of read peaks per kbp in naïve B-cells from ENCODE. **(B)** Correlation of gene depletion with potential sgRNA target sites. Log_2_ read frequency fold-change for day 0 to day 15 was binned by number of perfect or single base mismatches (within the first 12 bases of sgRNA) to the human genome.

For each feature, we also compared the union of the fourth and fifth bins (6,001-10,000) to the union of the last two bins (16,001-19,034). This comparison revealed that mutation rate and gene-trap score differences flattened out after the first 2,000 to 4,000 genes (p-values >0.05, two-tailed Wilcoxon test), whereas the other features still correlated with gene rank in the latter bins to varying degrees (PPI degree, p=0.00335; mRNA expression, p=0.0072, KBM7 score, p=3.94e-266; DNAse 1 hypersensitivity, p=1.27e-46). The tight concordance between the NALM-6 and KBM7 scores over all bins was expected given that the same sgRNAs were used to target all RefSeq genes included in both libraries. The strong correlation between rank score in the NALM-6 screen and DNA accessibility suggested a potential non-specific effect of sgRNA-directed DSB formation. The two additional significant correlations may be explained by the fact that mRNA expression level also correlated with DNA accessibility (Spearman’s rank correlation coefficient=0.34, p<2.2e-16), and protein interaction degree in turn correlated with gene expression (Pearson correlation coefficient=0.22, p<2.2e-16). These correlations are likely explained by unrepaired Cas9-mediated cleavage events in a fraction of cells leading to a DNA damage-dependent growth arrest and depletion from the pool independent of indel formation. Highly accessible DNA is more likely to be cleaved by Cas9 (Chari et al. 2015) and sgRNAs with multiple potential cleavage sites in amplified regions cause a greater fitness defect (Wang et al. 2015; Aguirre et al. 2016; Munoz et al. 2016), although this latter trend has not been observed in other screens (Tzelepis et al. 2016). To test this idea on a genome-wide scale we compared the predicted number of sgRNA matches in the genome to the level of depletion in the pool and indeed found a strong overall correlation (Fig. 2B). For this same reason, non-targeting control sgRNAs in the EKO library were actually enriched compared to the rest of the targeting sgRNAs in the pool (Fig. 1F), and therefore not an ideal control distribution. For subsequent analysis, we instead redefined the control set for RANKS as the 213,886 sgRNAs that targeted genes never reported as essential in any published screen, and we also removed all sgRNAs with more than two potential off-target cleavage sites and applied a fixed correction factor to sgRNAs with one or two potential off-target sites (see Supplemental Information and Fig. S4). This adjustment of the control sgRNA set allowed significance values to be reliably established, but left the relative gene rank order virtually unchanged (R-squared=0.978).

### Universal essential genes

The EKO library screen identified a total of 2,280 fitness genes in NALM-6 cells below a false discovery rate (FDR) of 0.05, which we generically termed essential genes (Table S3). Of these essential genes, 269 were specific to the NALM-6 screen: 225 corresponded to well-validated RefSeq genes, including 19 RefSeq genes not assessed previously, and 44 hypothetical genes annotated only in AceView or GenCode. The set of essential RefSeq genes in NALM-6 cells overlapped strongly with three previously identified sets of essential genes using different libraries, cell lines and methods (Fig. 3A) (Blomen et al. 2015; Hart et al. 2015; Wang et al. 2015). For direct comparative purposes between studies, we examined a set of 16,996 RefSeq genes shared between the EKO library and two published sgRNA libraries (Hart et al. 2015; Wang et al. 2015). We identified 486 genes that were uniformly essential across our NALM-6 screen and 9 previous screens in different cell lines (Hart et al. 2015; Wang et al. 2015), referred to here as universal essential (UE) genes (Fig. 3B; Table S3). This UE gene set was smaller than previously reported common essential genes in previous screens (Blomen et al. 2015; Hart et al. 2015; Wang et al. 2015) but was similarly enriched for processes required for cell proliferation and survival, including transcription, translation, energy metabolism, DNA replication and cell division (Fig. 3C).

**Figure 3.**
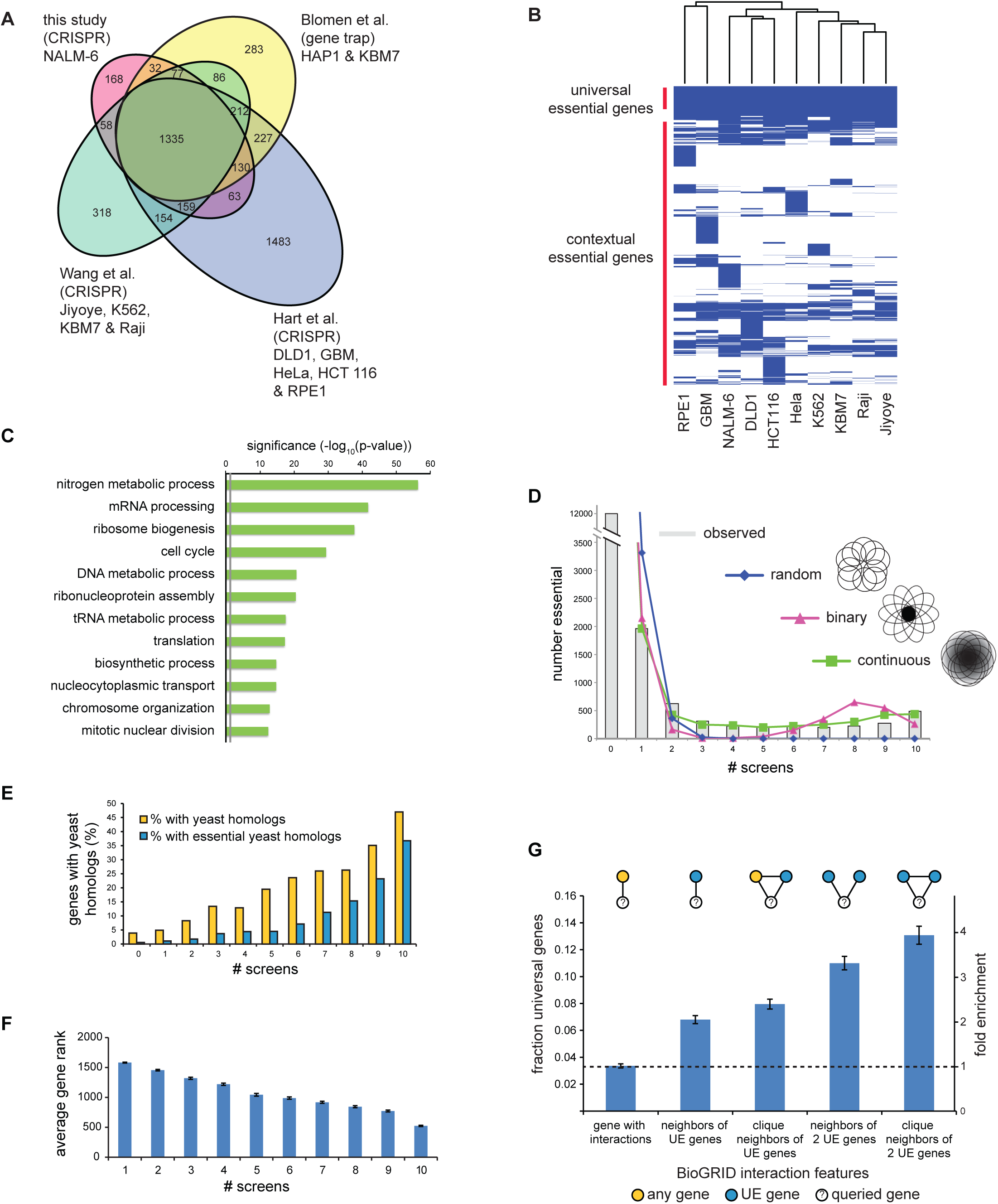
Universal essential genes. **(A)** Overlap between genes defined as essential in this study and three other studies (Blomen et al. 2015; Hart et al. 2015; Wang et al. 2015). The indicated overlap is between all essential genes identified in each study, i.e., an amalgam across all cell lines, in order to assess inter-library reproducibility. **(B)** Clustergram of essential genes across 10 different cell lines from CRISPR studies in panel A. **(C)** Biological processes enriched in UE genes. **(D)** Number of essential genes shared between cell lines. Experimentally observed essential genes are shown as a histogram. Indicated models to account for shared essential genes were fitted using maximum likelihood. See text for details. **(E)** Fraction of essential genes with orthologous non-essential or essential yeast genes as a function of cell line number. **(F)** Average score of each essential gene as a function of cell line number. **(G)** Fraction and relative enrichment of UE proteins in specific protein-protein interaction network motifs from BioGRID.

To investigate the nature of essential genes as a function of cell type, we plotted the number of shared essential genes across the 10 different cell lines. For clarity, we use the term contextually essential (CE) to refer to genes essential in more than one cell line but fewer than in all cell lines and lone essential (LE) to designate any gene that is uniquely essential in a single line. As opposed to a simple monotonic decay in shared essential genes as a function of cell line number, we observed a bimodal distribution whereby the number of shared essentials rapidly declined as the number of cell lines increased but then plateaued and peaked slightly at the maximum number of lines (Fig. 3D). To assess the potential effect of score threshold on the bimodal distribution, we examined the relative distribution of scores for essential versus non-essential genes as a function of cell line number (Fig. S5A). This analysis revealed that CE genes, especially those essential across many lines, often scored only slightly below the FDR threshold in other lines, consistent with genuine fitness defects that failed to reach significance in particular screens. However, it was also clear that many CE genes had no apparent fitness defect in particular cell lines, suggesting that bimodality was not merely driven by random effects in borderline essential genes (Fig. S5A). Indeed, we observed that bimodality was preserved over a wide range of essentiality thresholds (Fig. S5B). Variable essentiality across cell types may also reflect the expression of partially redundant paralogs (Wang et al. 2015). Consistently, genes with one or more close paralogs (>30% protein sequence identity) tended to have significantly lower essentiality scores in NALM-6 cells (p=3.6e-121, Wilcoxon test) and to be essential in fewer lines overall (p=1.5e-121, Wilcoxon test, Fig. S5C), but again this effect accounted for only a small fraction of the observed CE genes.

We assessed three different possible models that might help explain the observed bimodality of gene essentiality across cell lines. A random model represented the situation in which each gene was equally likely to be essential in any given cell line. A binary model corresponded to the scenario in which each gene was partitioned as either essential in all cell lines or not essential in any cell line, with an arbitrary constant for experimental noise. A continuous model represented the case in which each gene was assigned a specific probability of being essential in any given cell line (see Supplemental Information and Fig. S5D). The continuous model provided the best fit to the observed distribution as it was the only model that accounted for the prevalence of essential genes in the medial fraction of cell lines (Fig. 3D). This result suggested that gene essentiality is far from an all-or-none effect across different human cell types.

### Features of essential genes

Model organism studies have shown that essential protein coding genes are required for evolutionarily conserved processes in cell metabolism, macromolecular biosynthesis, proliferation and survival. Consistently, and as reported previously (Blomen et al. 2015; Hart et al. 2015), we observed that the more cell lines in which a gene was essential, the higher the probability that this gene possessed a budding yeast ortholog and that the ortholog was also essential in yeast (Fig. 3E) (Giaever et al. 2002; NCBIResourceCoordinators 2016). We also examined the converse question of why almost half of the essential genes in yeast were not universally essential in human cell lines. Out of 444 essential yeast genes with a human ortholog tested in all 10 cell lines, 387 orthologs were essential in at least one cell line, with a tendency to be essential in multiple cell lines. For the remaining 57 yeast genes that appeared to be non-essential in humans, 40 of these had GO terms linked to more specialized features of yeast biology. We also observed that as the number of cell lines in which a gene was essential increased, the greater the depletion of its sgRNAs from the library pool (Fig. 3F), such that UE genes were associated with significantly greater fitness defects. Consistent with the enrichment for crucial cellular functions, the set of GO terms associated with essential genes became progressively more restricted as essentiality spread over more cell lines (Fig. S6).

Proteins encoded by essential genes tend to cluster together within interaction networks in yeast (Hart et al. 2007), a feature also shown recently for human cells (Blomen et al. 2015; Wang et al. 2015). Using human protein interaction data from the BioGRID database (Chatr-Aryamontri et al. 2015), we observed that UE proteins tended to interact with each other more often than with random proteins (p<0.05, Fisher’s exact test) and associated preferentially within maximally connected subnetworks, referred to as cliques (for clique n=3, p<0.05, Fisher’s exact test) (Fig. 3G). These results suggested that clusters of UE genes carry out a limited set of indispensable cellular functions.

### Essentiality in human protein complexes

Essential genes tended to encode subunits of protein complexes as shown previously (Blomen et al. 2015; Hart et al. 2015; Wang et al. 2015), with a similar overall distribution as yeast essential genes (Fig. S7A,B; Table S4). We assessed the propensity of CE genes to interact with each other, and found that CE gene pairs essential in the same cell line were far more likely to encode subunits of the same protein complex than gene pairs that were essential in different cell lines (Fig. 4A). This result suggested that like UE genes, CE genes tended to encode essential modules in the proteome.

**Figure 4.**
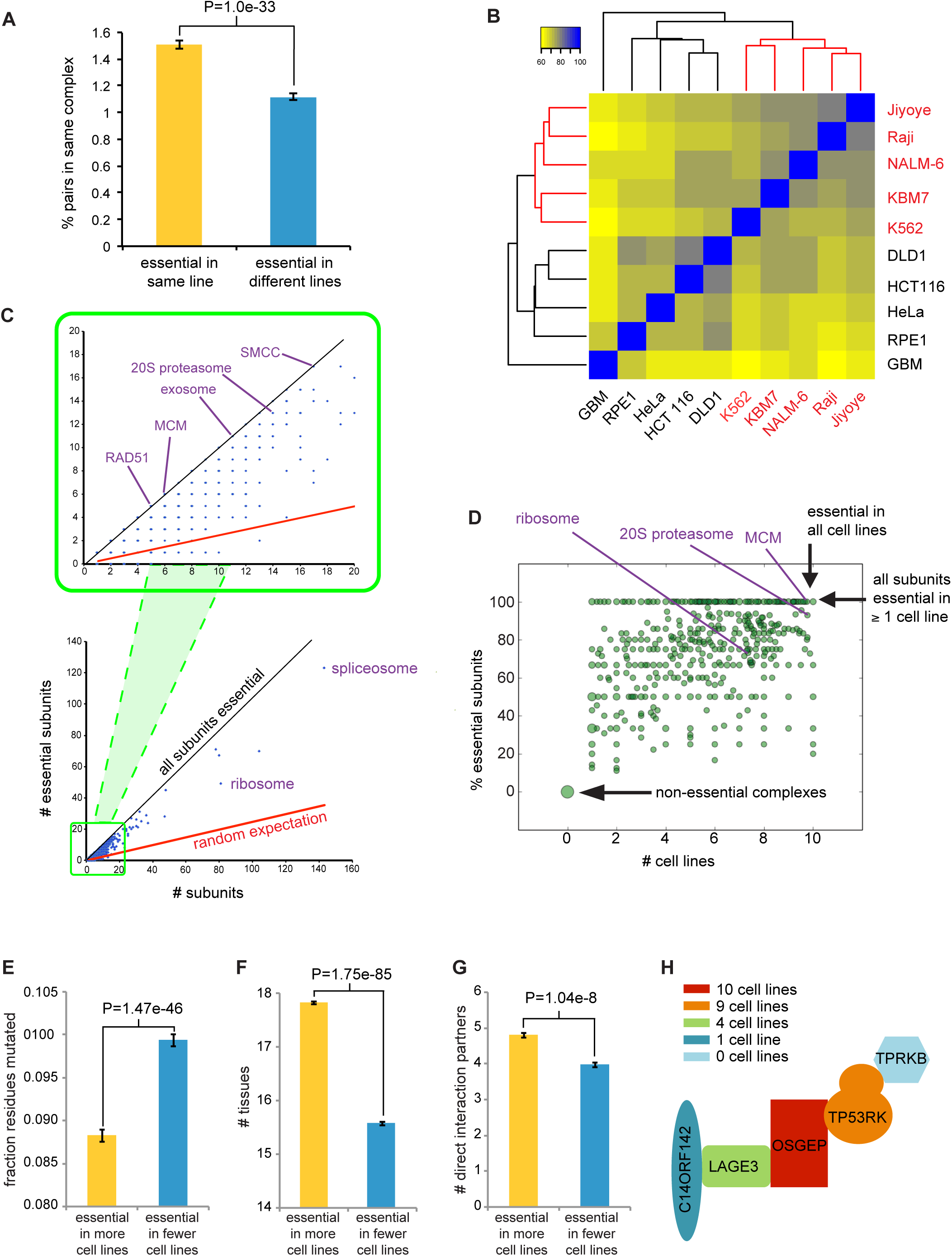
Essential subunits of protein complexes. **(A)** Pairs of essential genes in the same cell line tend to encode subunits of the same protein complex compared to essential genes unique to different cell lines. All possible pairs of cell lines were sampled by randomly choosing two pairs of genes per sample, one in which each gene was essential in line 1 but not in line 2 and then one additional gene essential in line 2 but not line 1. Probability of the first and second gene belonging to the same complex and that of the first and third gene were estimated from 10 million trials. P-value, Fisher’s exact test. Self-interacting proteins were excluded. **(B)** Cell lines clustered as function of shared essential protein complexes. All protein complexes were taken from the CORUM database release 17.02.2012 (Ruepp et al. 2010) and are listed in Table S4. **(c)** Distribution of essential subunits in protein complexes. **(D)** Fraction of essential subunits as a function of cell line number. Dot size is proportional to the number of complexes. **(E)** Mean fraction of mutated residues across 45 vertebrate species for all pairs of proteins that are part of a common CORUM complex and essential in at least one cell line. P-value, paired Wilcoxon rank-sum test. **(F)** Expression of subunits of CORUM complexes across 30 different tissues types as function of essentiality. **(G)** Proximity of subunits with at least 4 mapped subunits in a common PDB structure as a function of essentiality. **(H)** Variable essentiality of subunits of the KEOPS complex (Wan et al. 2016).

We predicted that shared essential genes should strongly cluster cell lines by cell type identity, but found that a number of cell lines did not segregate with similar lineages (Fig. S7C). However, when cell lines were clustered by shared essential complexes, defined as complexes with at least one essential subunit, the hematopoietic cell lines NALM-6, KBM7, Raji, Jiyoye and the colon cell lines DLD1 and HCT116 were all precisely grouped together (Fig. 4B; Fig. S7C,D). The functions carried out by essential complexes thus correlate with cell type identity more closely than the complete spectrum of essential genes.

At the level of individual complexes, a small number were comprised entirely of subunits essential in one or more cell lines, such as for the highly conserved SRB-and MED-containing cofactor complex (SMCC), exosome, the Rad51 homologous recombination repair complex and the DNA replicative helicase (MCM) complex (Fig. 4C). However, the vast majority of complexes contained a mixture of essential and non-essential (NE) subunits (Fig. 4D; Table S4). In order to assess whether the variation in essentiality between subunits of the same complex reflected the evolutionary history of the complex, we examined protein sequence conservation across 46 vertebrate species and found that subunits essential in more cell lines tended to be more conserved and expressed in more tissues than other subunits of the same complex (Fig. 4E, F). To test the notion that essentiality may reflect centrality in protein complex structure, we estimated the physical proximity of subunits for known complex structures in the PDB with at least four mapped subunits (Rose et al. 2015). Protein subunits that were essential in a greater number of cell lines tended to form more direct contacts with other subunits (Fig. 4G). As an example, the conserved KEOPS complex that mediates an essential tRNA modification reaction (Wan et al. 2016) contained a core catalytic subunit (Kae1) that was essential in all lines, a tightly linked core subunit (TP53RK) essential in nine lines, and three auxiliary subunits essential in four (LAGE3), one (C14ORF142) and zero (TPRKB) lines (Fig. 4H). The core Kae1-TP53RK subcomplex is flanked by the other subunits such that essentiality parallels structural centrality. The evolutionary plasticity of subunit essentiality is illustrated by the observation that Kae1 alone is sufficient for function in the mitochondrion, that three KEOPS subunits are essential in bacteria, and all five KEOPS subunits are essential in yeast (Wan et al. 2016).

### NALM-6-specific essential genes

Of all the genes tested in common between our study and nine previous CRISPR/Cas9 screens (Hart et al. 2015; Wang et al. 2015), 218 were uniquely essential to the NALM-6 screen. These LE genes had more protein interaction partners than NE genes (Fig. 5A, B) and exhibited higher expression levels in NALM-6 cells (Fig. 5A, C), two defining features of essential genes identified in other cell lines and model organisms (Blomen et al. 2015; Hart et al. 2015; Wang et al. 2015). However, LE proteins unique to NALM-6 cells were not as highly clustered in the protein interaction network as UE proteins (Fig. S8A, B), and had no more tendency to interact with UE proteins than NE proteins (Fig. S8C, D). Across all cell lines, while UE and CE proteins showed a strong propensity to interact, LE proteins were no more likely to interact with the UE core than NE proteins (Fig. 5D). These results suggested that LE proteins carry out a diversity of functions not strongly connected to the UE core, potentially as a consequence of synthetic lethal interactions with cell line-specific mutations.

**Figure 5.**
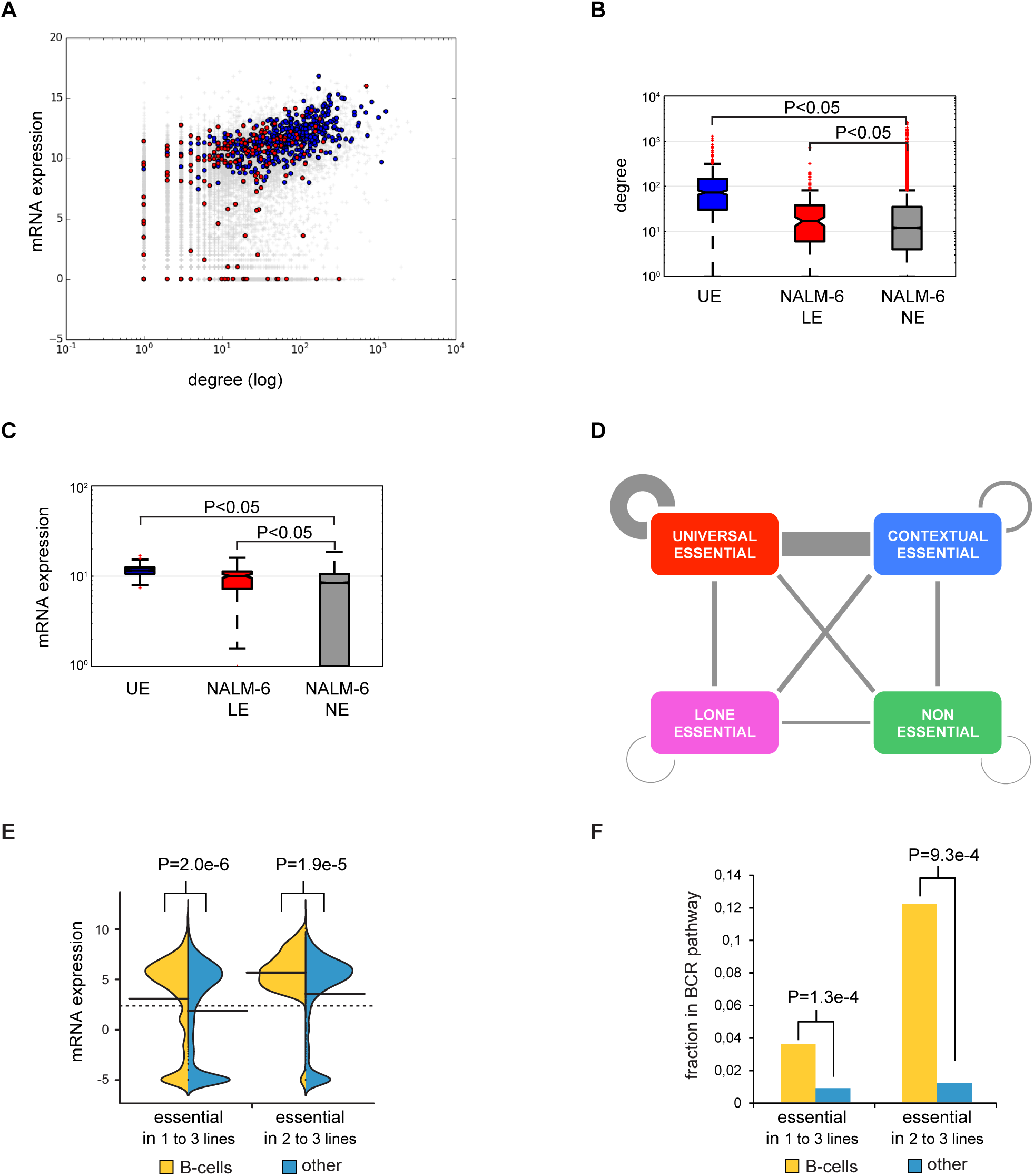
Cell type-specific essential genes. **(A)** Expression level as a function of protein interaction degree for essential genes unique to the NALM-6 screen (red) versus UE genes (blue). **(B)** Average interaction degree for proteins encoded by UE genes, LE genes in NALM-6 and NE genes. **(C)** Expression levels of UE genes, LE genes in NALM-6 and NE genes. **(D)** Interactions between UE, CE, LE and NE proteins. Counts were normalized by number of genes and interactions per class. **(E)** mRNA expression in NALM-6 cells of essential genes either unique to one or more B-cell-derived cell lines or unique to a randomly chosen equivalent number of other cell lines. **(F)** Participation in the B-cell receptor pathway (BCR) of essential genes either unique to one or more B-cell-derived cell lines or unique to a randomly chosen equivalent number of other cell lines.

The NALM-6 line bears an A146T mutation in NRAS (Bamford et al. 2004), analogous to mutations that activate KRAS in some leukemias (Tyner et al. 2009). As NALM-6 cells required the NRAS gene for optimal growth (Table S2), other NALM-6 specific essentials may be required to buffer the effects of oncogenic NRAS signaling. For example, two of the 11 components of cytochrome C oxidase (mitochondrial complex IV), COX6A1 and COX8A, were essential for survival exclusively in NALM-6 cells, as were two cytochrome C oxidase assembly factors, COA6 and COX16 (Table S3). Cytochrome C oxidase is known to be activated by oncogenic RAS and is required for survival of other cancer cell lines that bear an activated *RAS* allele (Telang et al. 2012).

In addition to NALM-6, the Raji and Jiyoye cell lines screened previously are each derived from the B-cell lineage (Wang et al. 2015). Surprisingly, of the 351 genes uniquely essential to one or more of these B cell lines (Table S2), only four genes were essential to all three lines, namely EBF1, CYB561A3, PAX5 and MANF. Two of these genes, PAX5 and EBF1 are key transcription factors that specify the B-cell lineage (Somasundaram et al. 2015), and were identified previously as shared essential genes between the Raji and Jiyoye cell lines (Wang et al. 2015). MANF, encodes mesencephalic astrocyte-derived neurotrophic factor and is expressed at high level in secretory tissues such as the pancreas and B-cells. MANF helps cells cope with high levels of protein folding stress in the endoplasmic reticulum (Lindahl et al. 2017) and also activates innate immune cells to facilitate tissue regeneration (Neves et al. 2016). CYB561A3, encodes a poorly characterized ascorbate-dependent cytochrome b561 family member implicated in transmembrane electron transfer and iron homeostasis (Asard et al.) but its role in B-cell proliferation or survival is not characterized.

We also examined all of the genes that were essential only in B-cell lines and found that these tended to be more highly expressed in NALM-6 cells than genes essential in an equivalent number of other cell lines (Fig. 5E), consistent with the higher expression of genes involved in B-cell proliferation in NALM-6 cells. These essential genes were significantly more likely to participate in the B-cell receptor (BCR) signaling pathway (Fig. 5F), which is often hyper-activated in chronic lymphocytic leukemia (Croft et al. 2011; Seda and Mraz 2015). Concordantly, disruption of this pathway reveals vulnerabilities specific to the B-cell lineage. For example, the BCR pathway components TSC2, PI3KCD, CD79B and CD19 were all essential in two of the three B-cell lines tested to date (Table S3).

### Residue-level features predict phenotypic effects of sgRNA-directed in-frame mutations

A fraction of indels introduced into genomic DNA following error-prone repair of a Cas9-mediated DSB will span a multiple of 3 base pairs such that the phenotypic effect will depend on the precise function of the mutated residue. For each sgRNA in the EKO library that targeted the 2,236 essential genes in NALM-6 cells, we identified the codon that would be subjected to Cas9 cleavage and hence the residue most likely to be affected by in-frame indels. We found that sgRNAs targeting predicted domain-coding regions were significantly more depleted from the pool than other sgRNAs targeting the same gene (p=1.8e-45, Fig. 6A). This result confirms a previous focused study on high density sgRNA-mediated targeting of ~200 genes (Shi et al. 2015) and generalizes the effect to the diverse classes of domains encoded by the entire genome. Based on the high degree of significance of this result, we asked whether other protein-level features would also correlate with sgRNA targeting sites. We found that sgRNAs targeting disordered protein regions were significantly less depleted than other sgRNAs targeting the same gene (Fig. 6B). The significance of this trend also held when the analysis was restricted to regions outside of Pfam domains (p=8.98e-6, data not shown). Disruption of more conserved regions caused significantly more depletion than for less conserved regions of the same gene (Fig. 6C). Targeted regions that encoded α-helices or β-sheets were also significantly more depleted than other regions (Fig. 6D). Buried residues within a protein structure were more depleted than accessible residues (Fig. 6E). Finally, sgRNAs targeting interfacial regions between two protein subunits were more depleted than non-interfacial regions of the same protein (Fig. 6F). Integration of each variable into a single linear multi-variate model, together with the number of potential off-target cleavage sites and predicted sgRNA efficiencies, revealed that every variable was significantly and independently correlated with relative sgRNA depletion (Table 1). This result indicated that each residue-level feature contributed to the phenotypic effects of in-frame mutations (Fig. 6G).

**Figure 6.**
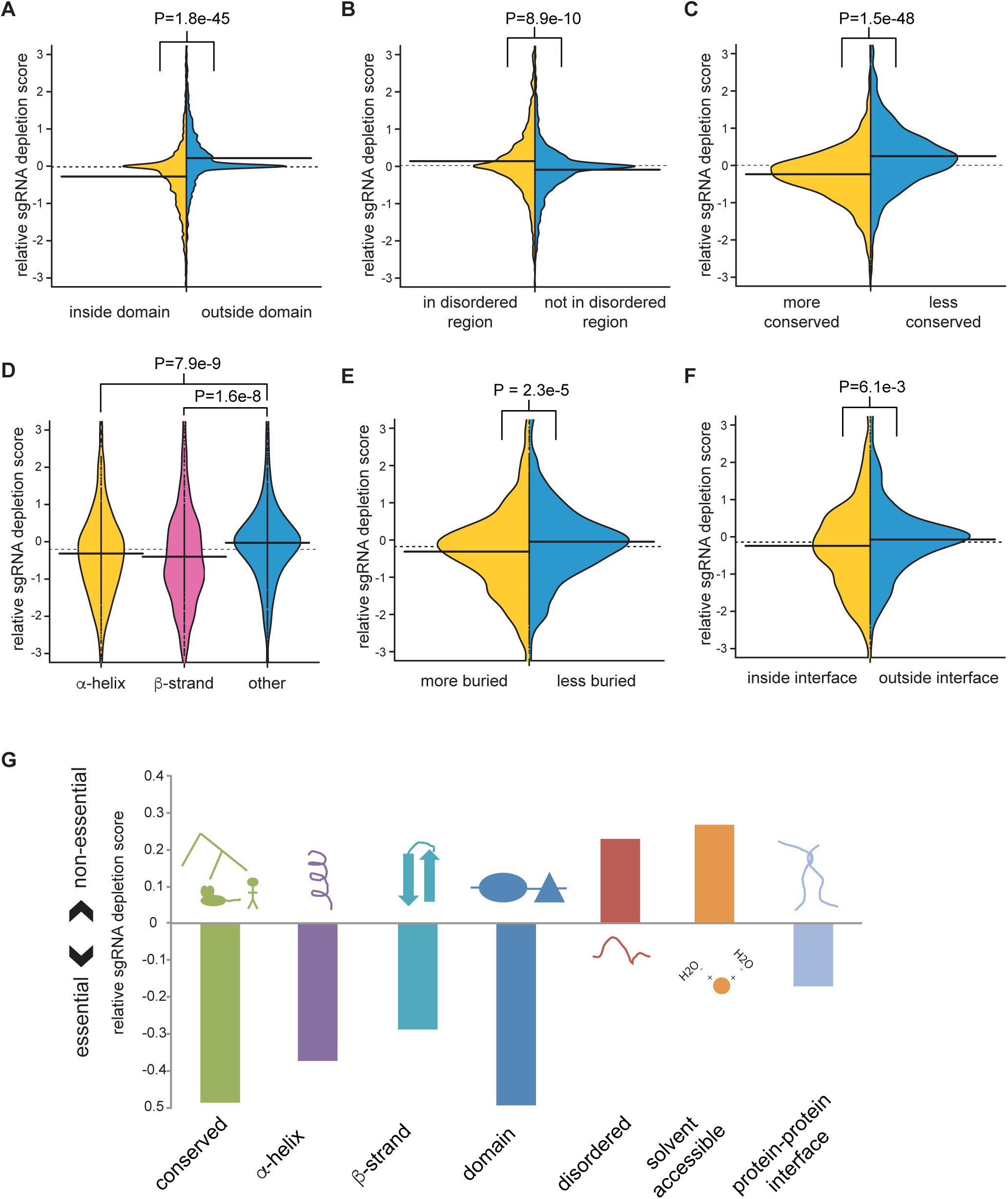
Correlation of sgRNA depletion with sub-gene and protein level features. Average log p-value of sgRNAs linked to each feature were normalized to the gene average for all sgRNAs. **(A)** Overlap with Pfam domains. **(B)** Predicted protein disorder of >40% probability as determined by UIPred (Dosztanyi et al. 2005). **(C)** Protein-level conservation of a 30 residue window, either more conserved or less conserved than gene average. **(D)** Residue burial, defined as the number of non-hydrogen atoms from non-adjacent residues present within 6 Å of the residue, for residues more or less buried than the average for the protein. **(E)** Residue proximity to a protein-protein interface. **(F)** Location of residues within an α-helix, an extended ²-strand or neither as annotated from PDB. **(G)** Summary of protein level features that influence sgRNA depletion from the library pool.

### Essentiality of alternatively-spliced exons

The EKO library was designed in part to target specific alternatively-spliced exons, many of which are found in essential gene loci. From the depletion of sgRNAs targeting these exons, we were able to classify individual exons within essential genes as either essential or non-essential (Table S5). We analyzed the 2,143 alternative exons within RefSeq-defined coding regions of essential genes in the NALM-6 screen, each of which was covered by at least 3 sgRNAs with ≥20 reads, to identify 462 exons with sgRNAs that were significantly depleted (FDR<0.05). When we compared these essential alternative exons to non-essential exons (N=592, FDR>0.3) in essential genes, we found that the essential exons were more likely to overlap protein domains (Fig. 7A), less likely to contain long disordered regions (Fig. 7B), more conserved across 46 vertebrate species (Fig. 7C), and more highly expressed at both the protein (Fig. 7D) and mRNA (Fig. 7E) level. Importantly, mRNA expression analysis showed that all 462 essential exons were expressed in NALM-6 cells. We also mapped the exons to full-length isoforms in the IsoFunct database, which assigns Gene Ontology functions to individual protein isoforms (Panwar et al. 2016). Isoforms that contained essential exons were more likely to be functional (i.e., higher IsoFunct scores) than isoforms with non-essential exons (Fig. 7F). The non-essential nature of particular exons may reflect structural features or protein interactions associated with the exon encoded region. For example, the anaphase-promoting complex/cyclosome (APC/C), contains a non-essential exon in the essential gene ANAPC5, which interacts with ANAPC15, itself a non-essential component of the complex (Fig. 7G) (Chang et al. 2015). These results suggested that the EKO library can effectively distinguish essential from non-essential alternatively-spliced exons within essential gene loci, and that many alternatively spliced exons of essential genes are non-essential.

**Figure 7.**
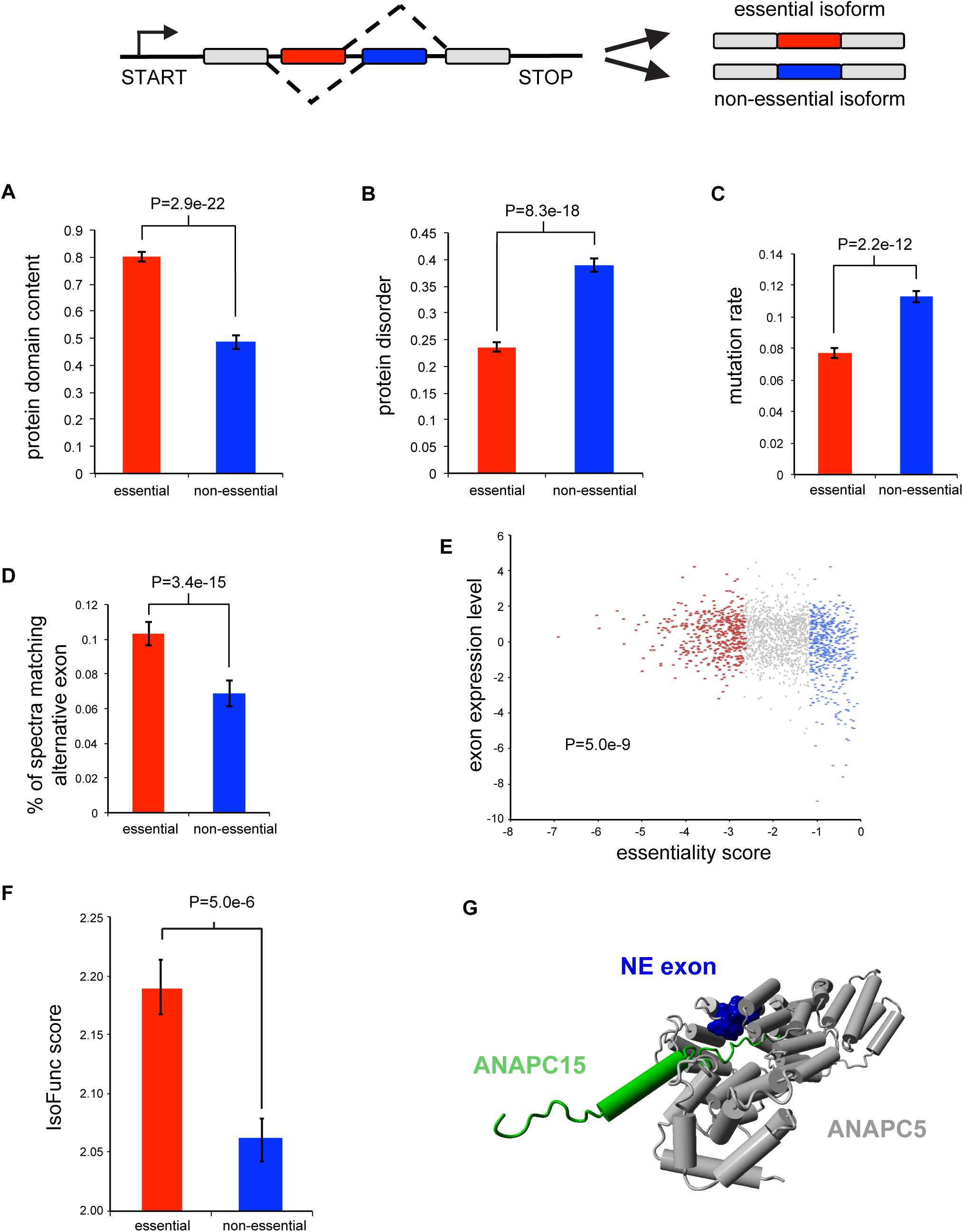
Alternatively spliced exons in essential genes. **(A)** Fraction of exons that overlap a Pfam domain (e-value <10^-5^) for essential exons (FDR<0.05) and non-essential exons (FDR>0.3) in essential genes. **(B)** Average IUPred predicted probability of exon residues belonging to a long disordered region. **(C)** Average fraction of residues mutated relative to human in 45 vertebrate species. **(D)** Average number of matched mass spectra per residue across human tissues. **(E)** Log_2_ RNA-seq read density in the exon normalized to gene average. **(F)** Average IsoFunc fold-change, representing the likeliness that a specific isoform codes for a given GO function for isoforms of essential genes. **(G)** Non-essential alternative exon (blue) in the essential gene ANAPC5, within the anaphase promoting complex interacts with ANAPC15, a non-essential component (green) of the complex.

### Genetic analysis of hypothetical genes

The EKO library was also designed to target unvalidated hypothetical genes that are currently absent from the RefSeq database (NCBIResourceCoordinators 2016). We identified these hypothetical genes from the AceView and GenCode databases, which annotate loci on the basis of expressed sequence tag (EST) and/or RNA-seq evidence. Because many hypothetical genes may be expressed pseudogenes present in more than one copy in the genome, we excluded all sgRNAs with close mismatches to the genome in order to avoid depletion effects due to multiple cleavage events. When we considered only sgRNAs with a single potential cleavage site, we identified 44 essential genes (FDR<0.05) that were absent from RefSeq (Table S3). Similar to most poorly annotated hypothetical genes, these genes tended to encode short polypeptides, with a median length of 153 residues and a maximum length of 581 residues (Fig. 8A). We found that these essential hypothetical genes were more highly expressed across a range of tissues than the 2,000 hypothetical genes with the least depleted sgRNAs (Fig. 8B). We also analyzed a comprehensive mass spectrometry-based proteomics dataset that covers 73 tissues and body fluids (Wilhelm et al. 2014) and found that the 500 hypothetical genes with the strongest sgRNA depletion scores were more likely to have evidence of protein expression than the 2,000 genes with the lowest sgRNA depletion scores (Fig. 8C). However, alignments across 46-vertebrate genomes revealed that the essential hypothetical genes were not more conserved than their non-essential counterparts (Fig. 8D), suggesting that the essential functions were acquired through recent evolution. These results suggest that at least a fraction of newly evolved uncharacterized hypothetical genes are likely to be expressed and perform important functions.

**Figure 8.**
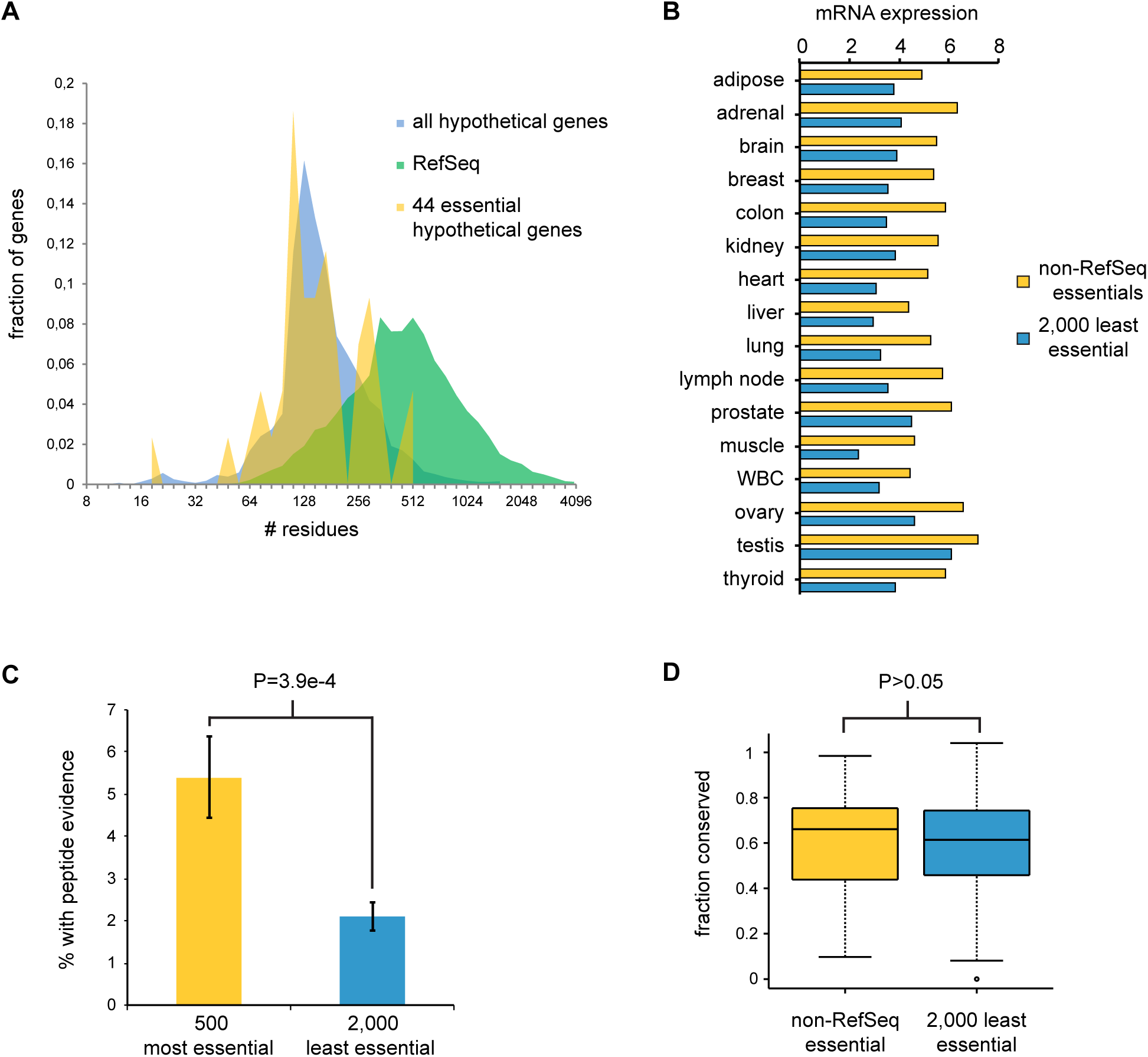
Correlated features of predicted hypothetical genes. **(A)** ORF length distribution of RefSeq genes, hypothetical ORFs from AceView or GenCode covered by the EKO library. The 44 hypothetical ORFs with significant (FDR<0.05) essentiality scores in the NALM-6 screen are indicated separately. For genes with multiple transcripts, only the length of the longest predicted protein was considered. **(B)** Mean log_2_(reads/kb) across 16 tissues from the Human BodyMap 2.0 for 44 essential and the 2,000 least essential hypothetical genes. **(C)** Fraction of genes with uniquely matched mass spectra at FDR<0.001 for the 500 most essential and the 2,000 least essential hypothetical genes. **(D)** Fraction of residues conserved across 46-vertebrate genomes for the 44 essential and the 2,000 least essential hypothetical genes.

## Discussion

Genome-wide collections of genetic reagents have allowed the identification of essential genes and insights into the functional architecture of model organisms. The first genome-wide CRISPR/Cas9 and gene-trap screens have defined a draft map of essential gene functions across a variety of human cell types (Blomen et al. 2015; Hart et al. 2015; Wang et al. 2015). Here, we have applied the high-complexity EKO library to define new essential features at the level of protein residues, alternatively spliced exons, previously uncharacterized hypothetical coding regions, and protein complexes.

### A continuum of gene essentiality

We combined our screen data with two previously published CRISPR/Cas9 screen datasets to define a minimal set of 486 UE genes across 10 different cell lines of diverse origins. As opposed to a simple monotonic convergence on a core set of essential genes, we observed an unexpected bimodal distribution of essential genes as a function of the number of lines screened. This distribution was best explained by a continuous variation in the probability of gene essentiality as opposed to a simple all-or-none binary model. This continuous probability model likely reflects genetic and/or epigenetic background effects, wherein different cell line contexts provide different levels of buffering against the loss of potentially essential genes. In this sense, the number of cell line backgrounds in which a gene is essential can be thought of as a form of genetic interaction degree, whereby the interaction occurs between any gene and a complex genetic background, as opposed to between two genes. In yeast, strain background markedly affects gene essentiality due to high order genetic interactions (Dowell et al. 2010), such that the essentialome of a particular cell must be defined in the context of a precise set of genetic and/or epigenetic parameters.

The essentialome may be thought of as an onion with multiple layers that become progressively more context-specific. As we show here, UE genes at the center of the onion make quantitatively greater fitness contributions than progressively more cell line-specific essential genes. The middle layers of the onion correspond to the plateau in the bimodal distribution in which genes are essential in specific cell lines. The definition of essentiality is obviously influenced by the definition of experimental thresholds, but the continuum of essentiality is nevertheless apparent regardless of the specific threshold. Based on the current but limited analysis, we estimate that this consensus essentialome for 8 or more cell lines will encompass approximately 1,000 genes. This number intriguingly is close to the number of 1,114 essential genes required for yeast growth under optimal growth conditions (Giaever et al. 2002).

It is important to note that this quantitative model of essentiality does not rule out the existence of a set of true UE genes. In fact, the continuum model predicts a smooth decline out to the maximal number of lines, in contrast to the sharp step increase observed experimentally at the maximum. This observation suggests the existence of a small set of UE genes that may be qualitatively distinct from the continuum of essentiality captured by the model. This set of essential functions likely corresponds to the structural and enzymatic core of the proteome, to which other essential functions have been added and subtracted through the course of evolution.

### A continuum of essential subunits in protein complexes

As in model organisms and as suggested by previous studies, the proteins encoded by UE genes tend to cluster together into modules in the protein-protein interaction network. This correlation between centrality and lethality is a robust feature of biological networks. Although originally posited to reflect the central location of essential proteins in network graphs (Jeong et al. 2001), this topological argument has been overturned in favor of the idea that essential proteins perform their functions as complexes (Zotenko et al. 2008). This notion has in turn been supported by the claim that protein complexes exhibit a tendency to be composed of mainly essential or mainly non-essential subunits (Hart et al. 2007; Ryan et al. 2013), such that the interactions of essential protein subunits would naturally cluster together. A surprising outcome of our analysis across multiple different human cell lines is the extent to which subunit essentiality for any given complex varies between cell lines. Our analysis of structural data suggests that UE subunits form the functional and structural core of essential protein complexes, to which CE, LE and NE subunits are appended. This observation probably reflects the fact that protein machines have acquired progressively more subunits through evolution, and that more recently evolved subunits tend to lie outside the essential structural core (Kim et al. 2006; Blomen et al. 2015). This variable essentiality of protein subunits is consistent with the lethality-centrality relationship (Fraser et al. 2002; Zotenko et al. 2008), but suggests that network evolution may also help drive the observed correlation (Coulombe-Huntington and Xia 2017).

### Cell line-specific essential genes and synthetic lethality

Only 4 essential genes were uniquely shared between the B-cell derived Raji, Jiyoye and NALM-6 lines, and only 35 genes were uniquely essential in B-cells and shared between two of the three lines. Similarly, a recent study identified only five essential genes shared between five acute myeloid leukemia (AML) cell lines, and 66 shared genes between at least three of the five lines (Tzelepis et al. 2016). These results suggest that genetic and epigenetic variation between lines may dominate the cell line-specific essentialome. Indeed, the majority of essential genes identified in human genome-wide screens to date are unique to single cell lines. In comparisons of wild type and laboratory yeast strains, which exhibit similar sequence variation as any two human individuals (Engel et al. 2016), strain-specific essential genes have a complex contextual basis due to multiple undefined genetic modifiers (Dowell et al. 2010). It seems likely that many cell type essential genes will reflect specific synthetic lethal interactions associated with the unique spectrum of cancer-associated mutations in any given cancer cell line. If on average each mutation yields 20 synthetic lethal interactions (Blomen et al. 2015; Costanzo et al. 2016), only 10 cell line-specific loss of function mutations would be needed to account for a specific essentiality profile of 200 LE genes. As an example, several components of cytochrome C oxidase and its assembly factors were specifically essential for survival in NALM-6 cells, potentially as a consequence of oncogenic RAS signaling (Telang et al. 2012), which is known to cooperatively trigger senescence in conjunction with mitochondrial defects (Nakamura et al. 2014). From a therapeutic perspective, the variable subunit essentiality of protein complexes suggests that a window of genetic sensitivity may exist for essential functions that are partially compromised in cancer cells (Hartwell et al. 1997; Hartman et al. 2001).

### Evolution of new functions

Recent RNA-seq and proteomics studies have illuminated the magnitude of alternative splicing in mammals and the attendant proteomic and phenotypic diversity generated by this mechanism (Barbosa-Morais et al. 2012). Different splice isoforms can have radically different functions, as for example pro-or anti-apoptotic splice variants of *BCL2* (Kontos and Scorilas 2012). Limited systematic studies on dozens to hundreds of isoforms suggest that exon composition can often dramatically alter protein interaction profiles (Ellis et al. 2012; Yang et al. 2016). Our genome-wide screen with the EKO library suggests that a large fraction of alternatively spliced exons in essential genes are not required for cell survival. This result suggests that alternative splicing has evolved as a means to diversify protein structure and interactions without compromising essential functions. Our data also shows that essential exons in general tend to encode structured protein domains that are more highly conserved and highly expressed, whereas non-essential exons tend to encode intrinsically disordered regions. Non-essential exons likely represent the first step towards the evolution of new functions, some of which may be then destined to become essential.

In contrast to the generation of new genes by duplication (Ohno 1970), the de novo appearance of new protein-coding genes is a poorly understood but nevertheless important evolutionary mechanism. For example, in mammals 1,828 known genes are unique to the primate lineage and 3,111 unique to rodents (Zhang et al. 2010). Proto-genes are genetic loci that produce very short proteins unique to each species and it has been suggested that a large pool of such potential genes exist in yeast (Carvunis et al. 2012). If a proto-gene assumes a beneficial function, it will be subjected to selective pressure and co-evolve across related species, such that even young genes can rapidly acquire essential functions (Chen et al. 2010). However, the detection of such proto-genes is hampered by the absence of sequence conservation and short length, which often precludes detection by mass spectrometry (Bruford et al. 2015). As shown here, CRISPR/Cas9-based screens allow the systematic functional analysis of hypothetical human genes. We detected an essential function for 44 hypothetical genes that was consistent with mRNA and protein expression data. These essential hypothetical gene functions were also evident across 36 different EKO library screens under various chemical stress conditions (data not shown). Although we cannot exclude the possibly that some of these loci may be non-coding RNA genes or pseudogenes, it is probable that these short reading frames represent newly evolved human genes. The biochemical interrogation of the corresponding proteins should help identify the function of these new candidate genes. Although we restricted our analysis to currently annotated hypothetical loci, it will be feasible to systematically query all short open reading frames in the human genome with dedicated sgRNA libraries.

### High resolution disruption of the human genome

Genome-wide CRISPR/Cas9-based screens have ushered in a new era of systems genetics in human cells (Shalem et al. 2015). The EKO library demonstrates the capacity of high resolution CRISPR/Cas9 screens to define gene essentiality across scales in the human proteome. Analogous deep coverage sgRNA libraries and variant CRISPR/Cas9 strategies also enable the precise mapping of regulatory and non-coding regions in the human genome (Wright and Sanjana 2016). Systematic functional analysis of these vast unexplored regions of the genome and proteome will provide insights into biological mechanisms and the evolution of phenotypic complexity in human cells.

## Methods

### Genome-wide sgRNA library

A set of 181,130 sgRNA sequences that target most RefSeq genes (~10 sgRNAs per gene) were as reported previously (Wang et al. 2014). Similar design rules were used to target additional genes from the latest RefSeq, Aceview and GENCODE releases (~10 sgRNAs per gene), as well as 20,852 alternatively-spliced exons derived from 8,744 genes (~3 sgRNA per exon). A set of 2,043 non-targeting sgRNA sequences with no detectable match to the human genome were randomly generated. The complete set of 278,754 sgRNAs was divided into three sub-library pools of 92,918 sgRNAs for array-based oligonucleotide synthesis as 60 mers (Custom Array), each containing 3-4 sgRNAs per gene, 1 sgRNA per alternative exon and 681 non-targeting sgRNAs. Each pool was amplified by PCR, cloned by Gibson assembly into the pLX-sgRNA plasmid (Wang et al. 2014), expanded in plasmid format and converted to a lentiviral pool by transfection into 293T cells (see Supplemental Information and Table S6).

### Screen for essential genes

A doxycycline-inducible Cas9 clonal cell line of NALM-6 was generated by infection with a Cas9-FLAG lentiviral construct and puromycin selection, followed by FACS sorting and immunoblot with anti-FLAG antibody to select a tightly regulated clone. The inducible Cas9 cell line was infected with pooled lentivirus libraries at MOI = 0.5 and 500 cells per sgRNA. After 6 days of blasticidin selection, 140 million cells were induced with doxycycline for 7 days, followed by periods of outgrowth in the absence of doxycycline. sgRNA sequences were recovered by PCR of genomic DNA, re-amplified with Illumina adapters, and sequenced on an Illumina HiSeq 2000 instrument.

### Scoring of gene essentiality

A RANKS score was calculated for every gene targeted by ≥4 sgRNAs represented by ≥20 reads in one sample. A single-tailed p-value for each sgRNA was obtained by comparing its ratio to that of the control sgRNAs. The RANKS score was determined as the average log_e_ p-value of the sgRNAs targeting the gene or exon. Gene/exon p-values were generated by comparing the RANKS score to a control distribution of an equal number of control sgRNAs for million samples with FDR correction (Benjamini and Hochberg 1995). The 2,043 non-targeting sgRNAs were used as controls in calculating the original RANKS score for scoring method comparisons and the binned rank analyses. To control for non-specific effects of Cas9 cleavage, the entire EKO library was used as a control after removing sgRNAs that targeted previously documented essential genes (Blomen et al. 2015; Hart et al. 2015; Wang et al. 2015) and sgRNAs with ≥2 predicted off-target cleavage sites. A fixed correction factor was applied to sgRNAs with 1 or 2 predicted mismatches. For the identification of essential non-RefSeq genes, only sgRNAs with unique matches to the genome were used to avoid potential confounding cleavage events at paralogous loci.

Further details of experimental, computational and statistical methods are provided in the Supplemental Information. All materials are available upon request and code for RANKS may be obtained at https://github.com/JCHuntington/RANKS.

## Acknowledgements

We thank Driss Boudeffa, Bobby-Joe Breitkreutz, Manon Lord, Jennifer Huber, Philippe Daoust and Alfredo Staffa for technical support, and Traver Hart, Jason Moffat, Luisa Izzi and other members of the Tyers laboratory for helpful discussions. JC-H was supported by a Canadian Institutes of Health Research (CIHR) postdoctoral fellowship, KB by Cole Foundation and Fonds de recherche du Québec Santé studentships, YX by a Canada Research Chair in Computational and Systems Biology, and MT by a Canada Research Chair in Systems and Synthetic Biology. Funded by grants from the Natural Sciences and Engineering Research Council of Canada (RGPIN-2014-03892 to YX), the CIHR (MOP-126129 to MT), the Canadian Cancer Society Research Institute (703906 to MT), the National Institutes of Health (R01RR024031 to MT), and by an award from the Ministère de l’enseignement supérieur, de la recherche, de la science et de la technologie du Québec through Génome Québec to MT.

### Author contributions

TB and JC-H designed the EKO library; TB built the EKO library and performed the screen with assistance from KB; JC-H developed the RANKS algorithm and performed all statistical analyses; AC-A performed protein interaction analyses; TB, JC-H, A-CA, YX and MT wrote the manuscript.

### Disclosure declaration

The authors declare no conflicts of interest.

